# Sexual Selection as a Mechanism of Evolutionary Information Preservation

**DOI:** 10.64898/2026.04.14.718539

**Authors:** Zeki Doruk Erden

## Abstract

Sexual selection is among the most pervasive and costly features of biological reproduction, reaching its greatest elaboration in lineages of greater complexity and capability—costs that signal a structural function not adequately accounted for by only the existing explanations focused on immediate adaptive benefits. I propose that this function is the long-term preservation of evolutionary information: through Fisherian co-evolution between female preferences and male traits, historically adaptive traits are sustained beyond the environmental pressures that originally favoured them and can be rapidly reactivated should those conditions recur, making sexually selecting lineages more efficient responders to fluctuating environments. I test this with an agent-based simulation in which male trait and female preference vectors co-evolve across alternating selective and neutral phases; across a wide range of conditions, any degree of female choosiness consistently preserves previously selected traits substantially better than random mating, with female preferences themselves forming a more durable archive than the expressed trait. The results carry implications for evolutionary theory and for the design of artificial life systems in which long-term retention of adaptive structure is a relevant property.

## Introduction

Sexual selection is a near-universal feature of sexually reproducing organisms, reaching its greatest elaboration and intensity precisely in those lineages that have achieved the highest degrees of morphological complexity and behavioral capability. It imposes substantial costs—reduced fecundity, energy expenditure, and in many cases elevated mortality risk—that grow more extreme with the elaborateness of the traits and preferences it drives. The scale and universality of these costs, concentrated among organisms of greater complexity, strongly suggest that sexual selection plays a structural role beyond local or immediate adaptive advantage. The central observation that demands explanation is not merely that sexual asymmetry exists, but that the lineages in which it is most pronounced are those that went on to become the most complex and capable.

Two well-established accounts of the evolutionary value of sexual reproduction are the Fisher-Muller hypothesis (Udomchatpitak, 2019), which emphasises the speed advantage of recombination in combining beneficial mutations, and the Red Queen hypothesis (Mueller, 2019), which emphasises genotypic novelty as a defence against rapidly co-evolving parasites. The sexual asymmetry between selecting and selected sexes can itself be modelled as a game-theoretic outcome of the two sexes acting as strategic agents with differing reproductive costs—an explanation with some force. Yet neither the recombination-focused accounts nor the game-theoretic framing addresses the deeper pattern: why it is specifically the asymmetric, sexually selecting lineages, rather than randomly mating ones, that tend to give rise to the more complex organisms. A more complete explanation must address not just why recombination or asymmetry is locally advantageous, but why sexual selection on this scale is associated with the emergence of greater long-term evolvability.

I propose that sexual selection serves a structural function in the evolutionary process: the long-term preservation of evolutionary information^1^ in the face of changing adaptive environments. Fisherian runaway co-evolution between traits and preferences is well established; what I argue is that this co-evolutionary structure may confer an underappreciated role at the wider evolutionary scale: sexually selecting lineages do not merely adapt to present conditions but accumulate a record of historically adaptive states. If a trait that was once under strong selection becomes relevant again—as environments fluctuate or recur—it can be already present or rapidly reactivated in a population that has retained it, rather than having to be re-evolved from negligible levels. This makes sexually selecting lineages more evolvable over the long run, not just better adapted in the short run. Once a preference is established, Fisherian runaway dynamics co-evolve preference and trait in a self-reinforcing loop that progressively decouples both from their original environmental basis, creating an internalised selection pressure that sustains the trait beyond its period of direct environmental relevance.

Understanding how reproductive mechanisms shape long-term evolutionary dynamics is also a central concern for artificial life research. Designing systems that can be steered toward desired long-term behaviors requires knowing how reproductive and selection mechanisms influence the path and persistence of adaptation. If sexual selection biases populations toward retaining historically adaptive information, this is a functionally important property for artificial life systems to account for—whether long-term retention is desired or not.

I operationalise this as trait retention: the capacity of a population to preserve a previously selected trait through a subsequent neutral phase in which the environment no longer favours it. I ask whether female choosiness *τ* —the strength of discrimination in mate selection—modulates this retention, and whether stronger sexual selection leads to longer persistence of past adaptive information.

Using an agent-based simulation, I find that across a wide range of population sizes, selection strengths, durations, and genetic parameters, populations with any degree of female choosiness systematically outperform random-mating populations in trait preservation. I further show that female preferences co-evolve with traits and are themselves retained during neutral phases, suggesting that the population’s preserved adaptive information is encoded redundantly in both the expressed phenotype and the latent preference.

## Related Work

Several artificial life platforms incorporate sexual reproduction. Misevic et al. (2006) showed that sexual reproduction reshapes the genetic architecture of digital organisms in AVIDA, with consequences for the speed and structure of adaptation. Yaeger’s PolyWorld (Yaeger, 1994) includes rudimentary mate selection. Standard evolutionary algorithm frameworks, developed primarily for optimisation (Pétrowski and Ben-Hamida, 2017), employ recombination as a genetic operator but treat selection as an exogenous fitness function. None of these systems model female preferences as evolved, heritable traits that generate the kind of co-evolutionary dynamics central to this study.

A number of computational models do represent female preferences as evolving properties. Adamson (2013) studied male life-history evolution under female choice; van Dijk et al. (2010) developed an individual-based quantitative genetic approach to sexual selection; Dreżewski (2018) modelled pair formation through agent-based simulation; and Mutoh et al. (2014) examined frequency-dependent sexual selection. These models illuminate important dynamics—signalling evolution, pair formation, equilibria—but are oriented toward understanding how traits or mating systems arise rather than toward their persistence across changing environments. The Sexy Son hypothesis (Weatherhead and Robertson, 1979) identifies an indirect fitness benefit from preferences in polygynous systems, but likewise does not address the preservation of adaptive information across environmental regime changes.

An analogous problem of retaining past adaptive structure arises in another domain of adaptive dynamics. In machine learning, and more broadly in systems trained by statistical optimisation—whether by gradient descent or by evolutionary algorithms, which share many structural properties as optimisation procedures—sequential training on changing objectives causes previously acquired capabilities to be lost, a phenomenon known as catastrophic forgetting (Van de Ven et al., 2024; Chen and Liu, 2022). This is not an incidental pathology but a structural consequence of optimisation on a changing signal, and it is shared in principle by evolutionary systems facing shifting selection pressures. I invoke this parallel not to equate the biological and computational phenomena, but as a dual motivator: it underlines why the retention of past adaptations in changing environments is a fundamental problem for any adaptive system, and it points to the practical relevance of understanding the mechanisms that address it—potentially providing insights not only for evolutionary theory but also for the design of artificial life systems where the balance between retention and adaptation can be deliberately controlled. This paper is not an AI or optimisation paper; the parallel is contextual.

To my knowledge, no prior computational study has specifically examined whether sexual selection preserves historically adaptive traits across environmental regime changes. This is the question I address.

### Hypothesis

The central claim of this paper is that sexual selection, operating through Fisherian runaway co-evolution, functions as a mechanism of evolutionary information preservation: it maintains the traces of historically adaptive states within the population even in the absence of continued external selection pressure. I develop this argument in three parts.

#### The inadequacy of immediate-benefit explanations

The prevalence and intensity of sexual selection across complex organisms demands a structural explanation. Its costs are not minor—reduced fecundity, energy expenditure on elaborate traits, and in many cases direct survival disadvantages—and they become more extreme, not less, in organisms of greater complexity. Explanations that appeal to immediate adaptive benefits— recombination speed, parasite resistance—account for why sexual reproduction is valuable, but they do not explain why it is so consistently accompanied by intense and asymmetric mate choice. The sexual asymmetry itself can be analysed as a strategic outcome of a two-sex game with differing reproductive costs, yielding stable asymmetric equilibria. But this framing, too, leaves the central observation unexplained: it is specifically the lineages where asymmetric sexual selection operates most intensely that tend to give rise to organisms of greater complexity. The causal question is not why complex organisms exhibit sexual selection, but why sexually selecting lineages gave rise to the complexity we observe. That question points toward a structural property of sexually selecting populations that makes them more evolvable in the long run—not merely better adapted in the short run. This hypothesis is complementary to, not contradictory with, the prior explanations.

#### The information preservation mechanism

The mechanism I propose is not a new one. Fisherian runaway co-evolution between female preferences and male traits is well-established as a theoretical and empirical phenomenon in the literature. What I argue is that this mechanism, viewed over extended evolutionary timescales, performs a function of evolutionary information preservation: it converts historically contingent external selection events into enduring internal selection pressures.

The selecting sex—typically females—cannot directly evaluate the underlying genetic quality or functional capacity of potential mates. It instead evaluates observable proxies: morphological signals, behavioral displays, or other traits that initially correlate with genuine fitness advantages in the current environment.

This proxy-based evaluation generates an initial selective pressure that mirrors the functional demands of the external environment. Once a preference for such a proxy is established, Fisherian runaway dynamics take over: males with the preferred trait are more reproductively successful, daughters inherit both the preference and the predisposition toward the trait, and the process amplifies progressively, decoupling both from their original environmental basis.

The result is an internalised selection pressure that maintains the historically selected trait even after the external pressure that originally established it has disappeared. The population carries, in the alignment between female preferences and male traits, a record of past selective environments. Traits that would otherwise decay when external selection is removed are preserved, not because they are currently demanded by the external world, but because the internal dynamics of mate choice continue to reward them. Critically, this preservation is not merely passive: if the environment shifts back toward conditions that again favour the retained trait, the population can respond rapidly— re-expressing or expanding a trait that was sustained rather than re-evolving one that was lost. This positions sexually selecting populations as more efficient responders to recurring or fluctuating selection pressures.

This places sexual selection in a different functional category from natural selection. Where natural selection tracks immediate environmental demands, sexual selection can stabilise historically significant traits, enabling the evolutionary trajectory to be path-dependent rather than purely reactive. At the level of abstraction, this is analogous to a form of evolutionary memory: just as memory systems protect past representations from being overwritten by new information, sexual selection protects past adaptive states from being erased by changing environments. The proposed mechanism thus extends the functional role of sexual selection beyond mate quality assessment: it is a population-level mechanism that makes sexually selecting lineages more evolvable in the long run, not merely better adapted in the short run.

#### Implications for biology and artificial life

This perspective is offered from outside biology proper. The author is not in a position to identify specific traits in real organisms preserved by this mechanism, nor to conduct the field and molecular analyses a direct empirical test would require. Nor, to my knowledge, has any inquiry been specifically directed at whether known sexually selected traits show preservation patterns attributable to this mechanism rather than to ongoing natural selection or other candidate explanations for sexually associated traits such as honest-signal display ornaments. This gap may partly reflect a timescale issue: the dynamics described here operate over hundreds to thousands of generations, and trait stabilisation through sexual selection may have concluded millions of years before those traits became objects of study, making the historical signal effectively invisible today. The theoretical proposal and computational results below nonetheless provide a basis for precisely such an inquiry, and if the proposed function is real, it implies a significant and underspecified contribution to the long-term evolvability of sexually selecting lineages. The computational results here do not by themselves constitute strong evidence for this claim in natural organisms, nor do they imply that information preservation is the primary or sole cause of the association between sexual selection intensity and organismal complexity. The proposed function is offered as a structural possibility meriting empirical scrutiny from expert biologists—not a conclusion a computational study from outside biology can establish.

For artificial life research, the implications are more immediately tractable. In artificial evolutionary systems the researcher has direct control over reproductive and selection mechanisms. Understanding how sexual selection modulates the retention of historically adaptive structure gives the designer a lever over the long-term behavior of the evolutionary process—toward greater retention, greater plasticity, or a balance between them. This matters for systems intended to operate in changing environments and for those in which past functional capacity may need to be recovered.

### Model

#### Individuals and genetic representation

Every individual carries two *n*-dimensional real-valued vectors. The trait vector **T** is phenotypically expressed in males and latent in females; the preference vector **P** is expressed in females and latent in males. Both sexes carry and transmit both vectors, enabling co-evolution across sexes. Population size is *N* per sex. Throughout this paper the selected sex is referred to as male and the selecting sex as female for clarity. This is a simplification: in nature the roles are not always so cleanly divided, and cases of reversed or symmetric choosiness exist; the terminology is adopted here solely for readability.

#### Natural selection

At each generation, male *i* survives with probability *σ*(*s*_*i*_), where *σ*(·) is the logistic function and

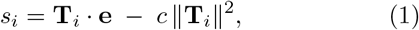

**e** ∈ ℝ^*n*^ is the environment vector, which encodes the direction and magnitude of the current selective pressure: fitness is maximised when the trait vector is aligned with **e** and decreases as the two diverge. *c* is a trait cost coefficient penalising large trait magnitudes. A minimum survival fraction *ρ* is enforced by promoting the highest-scoring remaining males if necessary, preventing population collapse.

#### Sexual selection and mate choice

Each female selects a mate from surviving males via a Boltzmann (softmax) distribution:

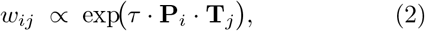

where *τ* ≥ 0 is the choosiness parameter. At *τ* = 0, mate choice is entirely random. As *τ* increases, females discriminate more strongly in favour of males whose trait vector aligns with their preference vector. Note that **P** does not enter the natural selection fitness function: preferences are internal evaluative structures rather than expressed physical traits, and as such they do not incur a direct survival cost analogous to the cost *c*∥**T**∥^2^ that penalises large male trait expression.

#### Inheritance

Each offspring inherits each vector component independently from one of its two parents with equal probability (Mendelian per-component inheritance), then receives an additive Gaussian mutation of standard deviation *σ*_*T*_ (traits) or *σ*_*P*_ (preferences). This preserves population-level genetic variance. Offspring are assigned a sex with equal probability and each sex is resampled to *N*.

#### Environmental regime and retention metrics

The environment follows a two-phase schedule. During the selection phase (*d*_*s*_ generations) the focal dimension *k* of **e** is set to the selection strength *s* and all other dimensions are set to zero, so that selection acts exclusively on component *k* of the trait. During the subsequent neutral phase (*d*_*n*_ generations) all dimensions of **e** are set to zero, removing all directional natural selection. Traits retain their cost *c* throughout both phases.

Retention is measured relative to the absolute value of the mean focal-trait component at the end of the selection phase, 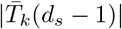 (here 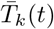 denotes the population mean of trait component *k* across the *N* males at generation *t*, and |·| its absolute value). Trials where this entry value is below 0.05 are excluded. Two complementary metrics are reported as median ± interquartile range (IQR) across 15 independent trials:

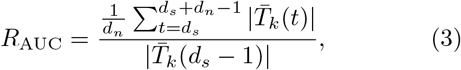

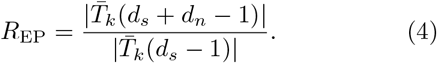

*R*_AUC_ (area-under-the-curve retention) measures average preservation over the entire neutral period; *R*_EP_ (end-point retention) measures the state of the population at the very end of the neutral phase. Both are computed analogously for female preferences 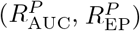. The two metrics behave differently depending on choosiness level. At *τ* = 0 the trait decays monotonically during the neutral phase, so the time-average (AUC) exceeds the terminal value: the AUC captures the early still-elevated portion of the decay and is therefore higher than the end-point. At *τ >* 0, Fisherian co-evolutionary dynamics can continue to amplify both trait and preference during the neutral phase—the internal mate-choice pressure persists as long as the co-evolutionary correlation is maintained—so end-point retention can meet or exceed AUC retention, and in several conditions exceeds 1.0. Values above 1.0 reflect ongoing Fisherian amplification beyond the selection-phase peak rather than mere preservation; sexual selection can sustain and expand the trait even in the complete absence of external selection. The metrics are complementary: AUC captures average retention across the neutral period; end-point captures the state at its close.

## Results

### Baseline: sexual selection preserves past traits

I first establish the effect under a baseline configuration: *N* = 200, *d*_*s*_ = 200, *d*_*n*_ = 200, *s* = 1.5, *n* = 3, *c* = 0.05, *σ*_*T*_ = 0.05, *σ*_*P*_ = 0.10, *ρ* = 0.30. I sweep *τ* ∈ {0, 0.5, 1, 2, 3, 5} with 15 trials per value using deterministic seeds.

Figure 1 shows the median trait trajectories. At *τ* = 0 the trait decays steadily once the neutral phase begins. At *τ >* 0 the trait persists at elevated levels, with the magnitude of persistence broadly increasing with choosiness. Table 1 provides quantitative detail. At *τ* = 0, median AUC retention is 0.705 and end-point retention is 0.391. Every *τ >* 0 condition substantially outperforms this baseline. AUC retention ranges from 0.766 to 0.960 and end-point retention from 0.818 to 1.722; several conditions achieve or exceed *R* = 1.0, meaning the trait is maintained at or above its selection-phase level. These results support the core hypothesis: sexual selection actively preserves previously acquired adaptive information, keeping traits available for rapid re-expression should the selective environment return.

**Table 1:**
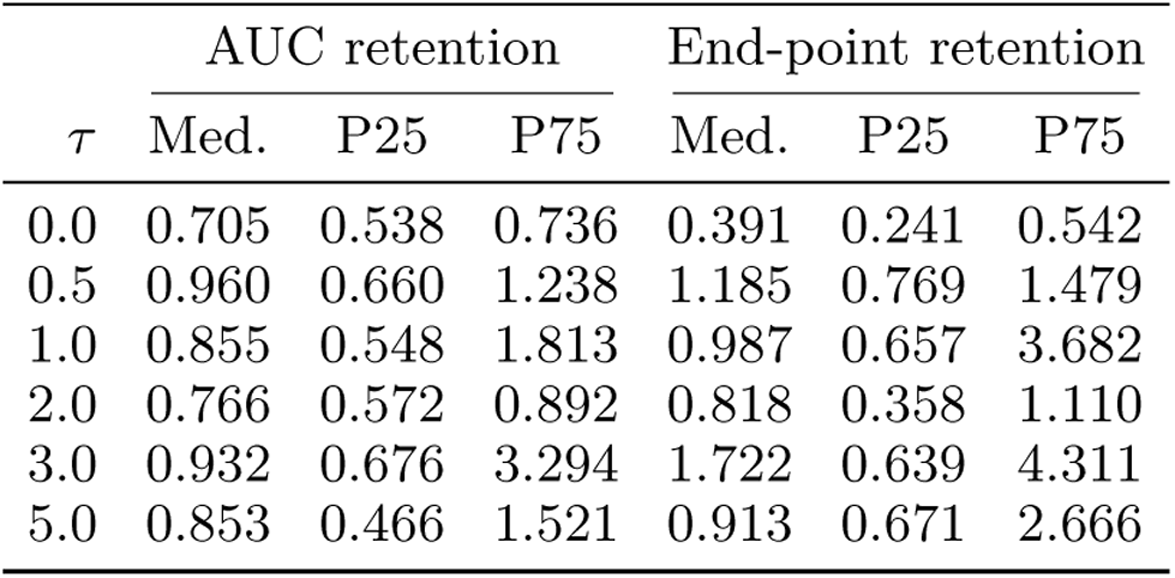
Baseline trait retention (AUC and end-point) per *τ*. *N* = 200, *d*_*s*_ = *d*_*n*_ = 200, *s* = 1.5. Median, P25, P75 across 15 trials.

**Figure 1.**
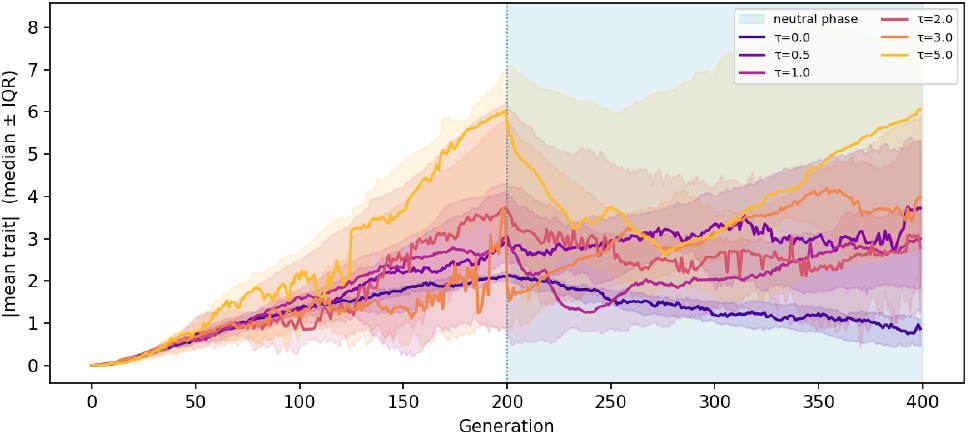
Median male trait 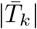 trajectories under the baseline parameter configuration, one line per choosiness level *τ*. Shaded bands indicate interquartile range across 15 trials. Blue region = neutral phase (*d*_*n*_ = 200 generations). At *τ* = 0 the trait decays steadily; at *τ >* 0 sexual selection maintains it above the randommating baseline.

**Figure 2.**
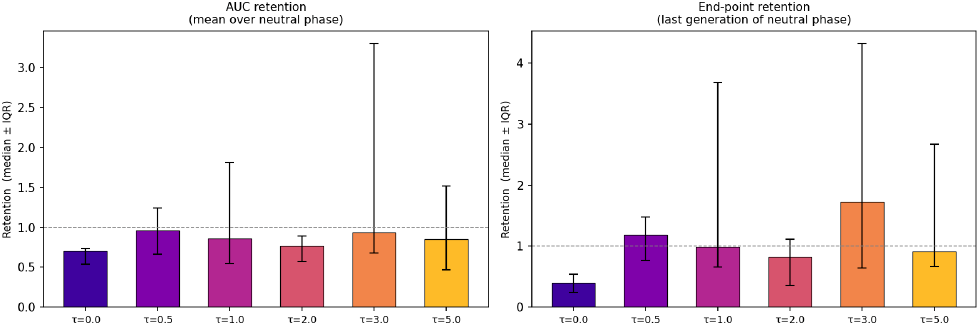
AUC retention (left) and end-point retention (right) per *τ* under the baseline configuration, median ± IQR across 15 trials. Dashed line at *R* = 1 indicates full retention. Any *τ >* 0 consistently outperforms the random-mating (*τ* = 0) baseline.

The gradient across *τ* levels is broadly positive but not strictly monotone, and inter-trial variability is high, particularly at large *τ* where runaway dynamics introduce instability. The primary finding is the robust binary distinction between *τ* = 0 and *τ >* 0: any degree of female choosiness provides a substantial information preservation advantage over random mating. End-point retention differences are statistically significant across all *τ >* 0 levels (Mann-Whitney *U, p <* 0.05); AUC differences reach significance at *τ* = 0.5 and *τ* = 3.0, with other *τ >* 0 values showing consistent directional advantages below threshold. Detailed statistical results are presented in the appendix.

### Robustness across parameter conditions

I tested the generality of the *τ >* 0 preservation advantage across diverse parameter settings: population size (*N* ∈ {50, 200}), regime duration (*d*_*s*_ = *d*_*n*_ ∈ {30, 200}), selection strength (*s* ∈ {0.3, 1.5}), trait dimensionality (*n* ∈ {1, 2, 3, 5, 8, 10}), trait cost (*c* ∈ {0.01, 0.02, 0.05, 0.10, 0.20, 0.50}), minimum survival fraction (*ρ* ∈ {0.05, 0.10, 0.20, 0.30, 0.50, 0.70}), and mutation rates (*σ*_*T*_ across five levels from 0.01 to 0.20, with *σ*_*P*_ = 2*σ*_*T*_). Table 2 summarises selected conditions, showing the *τ* = 0 baseline against the best-performing *τ >* 0 value. Detailed figures for all parameter sweeps are provided in the Appendix.

**Table 2:** Parameter robustness summary. For each condition, *τ* = 0 retention is compared with the best single *τ >* 0 value (shown in parentheses). † At *n* = 10, *τ* = 0 achieves the highest AUC of all values; the best *τ >* 0 is shown for completeness.

| Condition | AUC |  | End-point |  |
| --- | --- | --- | --- | --- |
| | $\tau=0$ | Best $\tau>0$ | $\tau=0$ | Best $\tau>0$ |
| Baseline | 0.705 | 0.960 (0.5) | 0.391 | 1.722 (3.0) |
| $N = 50$ | 0.803 | 1.133 (1.0) | 0.586 | 1.342 (0.5) |
| $d_s = d_n = 30$ | 0.963 | 1.306 (1.0) | 0.864 | 1.860 (3.0) |
| $s = 0.3$ | 0.568 | 2.227 (0.5) | 0.363 | 2.582 (0.5) |
| $n = 1$ | 0.650 | 1.230 (0.5) | 0.310 | 1.750 (1.0) |
| $n = 10^\dagger$ | 0.640 | 0.590 (3.0) | 0.380 | 0.630 (0.5) |
| $c = 0.01$ | 0.940 | 1.590 (1.0) | 0.880 | 2.100 (1.0) |
| $c = 0.50$ | 0.200 | 0.670 (3.0) | 0.050 | 0.860 (3.0) |
| $\rho = 0.05$ | 0.700 | 1.100 (0.5) | 0.390 | 1.170 (0.5) |
| $\rho = 0.70$ | 0.150 | 1.070 (3.0) | 0.010 | 1.490 (5.0) |
| $\sigma_T = 0.03$ | 0.750 | 2.710 (2.0) | 0.570 | 3.410 (2.0) |
| $\sigma_T = 0.20$ | 0.140 | 0.390 (1.0) | 0.030 | 0.270 (1.0) |

A consistent result across all tested conditions: at least one *τ >* 0 value outperforms *τ* = 0 in at least one metric. The gradient is sharpest under the baseline configuration. Several conditions degrade the gradient in informative ways, with some *τ >* 0 values underperforming—either because they are too weak to establish a co-evolutionary lock within the available time, or because higher values trigger interfering dynamics—yet even in these cases, some intermediate level of choosiness provides a measurable retention advantage over random mating.

Small population (*N* = 50): genetic drift dominates, producing high inter-trial variance. The *τ >* 0 advantage is detectable but unreliable.

Short regimes (*d*_*s*_ = *d*_*n*_ = 30): insufficient time for the co-evolutionary lock between **T** and **P** to form. Retention ratios are noisy and the gradient weak.

High dimensionality (*n* ≥ 8): two mechanisms degrade the gradient. First, preference dilution: only one preference component (*P*_0_) is reinforced by co-evolution with the selected trait, while the remaining *n* − 1 drift under mutation and contribute noise to mate-choice decisions, reducing the effective signal-to-noise ratio of female discrimination approximately as 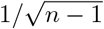. Second, spurious Fisherian runaway: drifting unselected preference dimensions can initiate their own co-evolutionary loops with the corresponding trait dimensions, redirecting selective energy away from the informative dimension. At *n* = 10, the AUC gradient collapses and *τ* = 0 outperforms all *τ >* 0 values, though the end-point metric still shows a modest *τ >* 0 advantage.

High cost (*c* = 0.50): natural selection actively opposes the trait in the neutral phase, narrowing the window in which sexual selection can counteract it. The *τ >* 0 advantage persists but is compressed.

These failure modes are informative: the preservation effect is most robust when populations are large enough to limit drift, evolutionary time is sufficient for co-evolution to establish, preferences are focused on few dimensions, and natural selection pressure against the trait is moderate rather than extreme. Across all conditions, however, some positive level of choosiness consistently provides more information preservation than none—the question is only which *τ* value and under what conditions the advantage is large enough to be clearly visible.

### Female preference co-evolution and retention

The analyses above concern the male trait. I now ask whether female preferences themselves co-evolve and are retained, using the same metric framework with 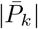 in place of 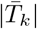.

Figure 3 shows parallel trajectories for 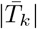 and 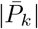 under the baseline parameters. At *τ* = 0, preferences remain near zero throughout: without directional mate choice, no selective pressure acts on **P**, and it drifts under mutation alone. At *τ >* 0, the Fisherian co-evolutionary feedback drives **P** upward alongside **T** during the selection phase, and both are maintained into the neutral phase.

**Figure 3.**
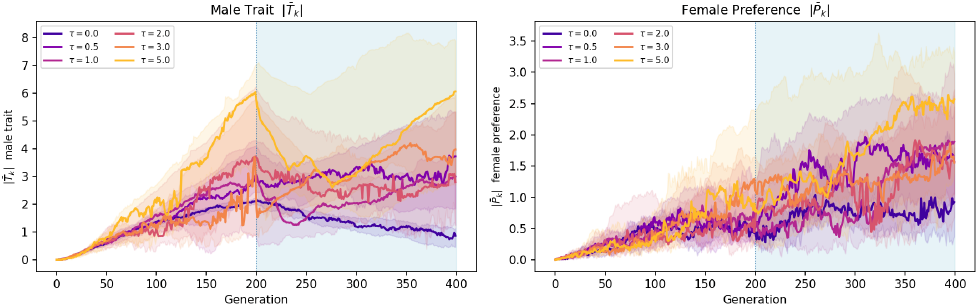
Median trajectories of male trait 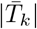 (left) and female preference 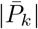 (right) under the baseline configuration. At *τ* = 0, preferences drift near zero and never co-evolve with the trait. At *τ >* 0, both rise together during the selection phase and are retained into the neutral phase (blue region).

Quantitatively, preference retention substantially exceeds trait retention at all *τ >* 0 levels. Median AUC preference retention ranges from 1.22 to 2.03, and median end-point retention from 1.55 to 2.69—well above the corresponding trait values and, notably, above 1.0 throughout, meaning preferences are not merely preserved but continue to increase during the neutral phase. Two factors account for this. First, unlike the trait, the preference incurs no natural selection cost: the fitness penalty *c* ∥**T**∥^2^ acts only on male trait expression, since preferences are internal evaluative structures with no direct physical expression that could be penalised by survival selection. Second, the Fisherian feedback can continue to amplify preferences in the neutral phase as long as some residual trait expression persists.

Figure 4 presents a direct side-by-side comparison of trait and preference retention per *τ*. The asymmetry is consistent: preferences form a more durable component of the stored adaptive information than the trait phenotype itself. This has a concrete functional implication for the reactivation advantage described above: when the environment shifts back toward conditions that favour the retained trait, the population benefits not only from residual trait expression but from a preference structure that is still actively directing sexual selection toward that trait—accelerating the re-expression of the historically adaptive phenotype.

**Figure 4.**
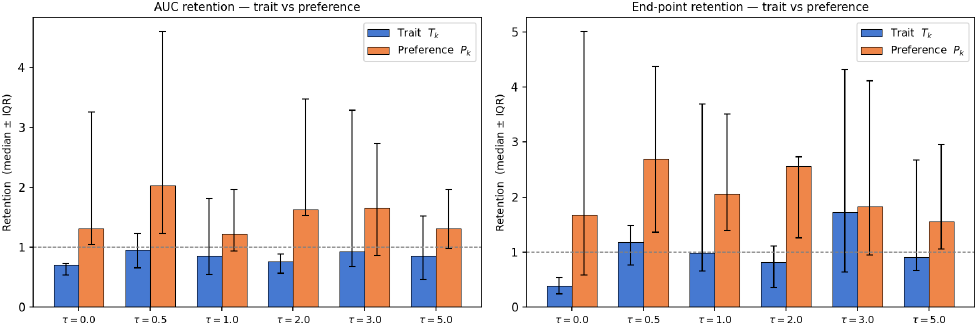
Direct comparison of male trait retention *R*_*T*_ (blue) and female preference retention *R*^*P*^ (orange) per *τ*, for AUC (left) and end-point (right) metrics, under the baseline configuration. Error bars indicate IQR. Preferences are consistently retained more strongly than traits, reflecting the absence of a natural-selection cost on the preference in the neutral phase.

## Discussion

### Sexual selection as an information preservation mechanism

The co-evolved (**T, P**) correlation created during a selection phase functions as an encoding of past environmental information: it specifies which traits were fitness-relevant in a previous regime and sustains a selection pressure toward their expression even after the regime changes. This creates path-dependence in the evolutionary trajectory: what a population has been selected for in the past shapes its current selective landscape through an intrinsic rather than extrinsic mechanism. The practical consequence is an asymmetry in re-adaptation: when a previously encountered environment returns, a population that has retained the trait— and especially the preference structure directing selection toward it—can re-express the adaptive phenotype far faster than one that must re-evolve it from negligible expression. A three-phase reactivation experiment—in which the selective environment is restored after the neutral phase—confirms this advantage directly; results and discussion are presented in the appendix.

The robustness of the *τ* = 0 versus *τ >* 0 distinction across diverse parameter settings is notable. The gradient across *τ* levels may require favourable conditions to emerge clearly, but the baseline contrast—any mate choice versus none—is detectable in nearly every tested configuration. This suggests that the information preservation effect is not a fine-tuned consequence of particular parameter values but a broadly operative consequence of the co-evolutionary feedback structure.

### Distributed and layered information storage

The preference results reveal that the population’s preserved record of past selection is distributed across two components: the expressed male phenotype **T** and the latent female preference **P**. The trait is subject to natural selection throughout: it is directly penalised by the cost *c* when the environment no longer rewards it. The preference is not: it is sustained by the co-evolutionary feedback alone and decays only through mutation. The preference therefore acts as a more durable archival layer, while the trait is the more dynamic, environmentally sensitive expression.

This asymmetry has a functional implication: should the environment become selective again for the historically favoured trait, populations with high *τ* are doubly advantaged. They retain not only residual trait expression but also the preference structure that directed selection toward that trait—enabling faster re-adaptation than a population that must rebuild both from scratch.

### Connection to adaptive optimization frameworks

Adaptive optimization systems—gradient-based machine learning and evolutionary algorithms—share a structural challenge: sequential optimisation on changing objectives tends to degrade previously acquired solutions. In machine learning this is studied as catastrophic forgetting (Van de Ven et al., 2024; Chen and Liu, 2022), where training on new tasks overwrites representations from prior ones; the same problem arises in evolutionary systems facing shifting selection pressures. Sexual selection provides an emergent biological analogue: the co-evolutionary lock between trait and preference protects against the erasure pressure of a changing environment. This raises the question of whether analogous mechanisms—internal selective structures that preserve historically useful representations—could be deliberately built into artificial evolutionary or learning systems for similar retention. This is relevant not only for optimisation and learning systems but for open-ended evolutionary systems intended to accumulate complex adaptive structure over extended timescales, where the capacity to retain historically built structure while continuing to evolve may be as important as adapting to immediate conditions.

### Broader methodological implications

The present work is an instance of a broader research approach: examining phenomena in evolutionary history from a functional and structural perspective to understand their influence on evolutionary dynamics. Even if such a property was not the dominant causal factor historically, once identified it can be incorporated into the design of artificial systems. The present results suggest preference co-evolution is one such property: a mechanism that, by its structure, preserves historically useful representations across changes in the selective environment. This points toward a broader program of inquiry: systematically examining features of evolutionary dynamics—reproductive mechanisms, population structure, developmental constraints—from this functional perspective, asking in each case what structural role they play in the long-term evolution of complexity and evolvability, and whether that role can be replicated or controlled in artificial evolutionary and artificial life systems.

### Limitations

The model is deliberately minimal. Trait and preference vectors are continuous and haploid; genetic architecture is non-epistatic; the environment vector is fixed and axis-aligned; regime changes are abrupt; and female costs of choosiness are not modelled (an experiment exploring the effect of such costs is presented in the appendix). More realistic settings—diploid inheritance, partial or gradual environmental change, multi-objective selection, rotating selection axes—would clarify the generality and the quantitative limits of the effect. The high dimensionality results additionally suggest that the fidelity of sexual selection as an information preservation mechanism depends critically on the degree to which preferences are focused on ecologically informative dimensions, a constraint that biological systems may enforce through developmental or sensory bottlenecks. Finally, in the present model, the selected sex (male) expresses only traits and the selecting sex (female) expresses only preferences: neither sex evolves both. This is an intentional simplification to keep the dynamics tractable and interpretable, but a more complete model would allow both sexes to carry and express both traits and preferences, with their relative magnitudes determined by the dynamics of the system rather than fixed by assumption. Relaxing this asymmetry is an important direction for future work.

## Conclusion

I have shown computationally that female choosiness acts as a tunable mechanism for the long-term preservation of evolutionary information. Populations with any degree of mate discrimination consistently retain previously selected traits better than random-mating populations across a wide range of ecological and genetic conditions. Female preferences co-evolve with traits and are preserved more durably than the traits themselves, distributing the population’s record of past selection across both phenotype and preference.

From this perspective, sex and sexual selection can be understood not merely as solutions to immediate adaptive problems—recombination efficiency, parasite resistance—but as a structural mechanism that makes the evolutionary process path-dependent and informationally persistent. The value of this mechanism, like the value of any information preservation system, lies not only in the present moment but in the system’s capacity to respond efficiently when past conditions reassert themselves—re-expressing traits that were retained rather than re-evolving traits that were lost.

This view opens several directions for future inquiry. Do real organisms exhibiting traits associated with strong historical sexual selection show evidence of environmental retention not explainable by natural selection alone? Can artificial life systems with designed preference co-evolution outperform standard evolutionary algorithms in cyclically changing environments? And how do preference dimensionality and cost structure modulate the fidelity of this information preservation function? More broadly, the results suggest that mechanisms analogous to sexual selection may be worth incorporating in artificial adaptive systems where long-term information retention is valued alongside immediate performance—and that studying the structural and functional roles of evolutionary phenomena more generally is a productive path toward understanding, and potentially designing, systems with greater long-term evolvability.

## Acknowledgements

No funding was received for this work. AI tools (Claude by Anthropic) were used to assist with code generation and text editing.

## Footnotes

1 By evolutionary information I mean the record, carried in a population’s heritable composition, of which trait configurations were favoured by the selective environments it has experienced.

